# Migratory connectivity of pelagic predators between tropical Atlantic islands and seamounts revealed through passive acoustic telemetry

**DOI:** 10.64898/2026.06.13.732048

**Authors:** SB Weber, AS Afonso, L Clarke, DJ Curnick, M Cranfield, L Henry, K Jones, J Letori, P Oliveira, D Simpson, T Simpson, D Viana, NE Hussey

## Abstract

**BACKGROUND:** Oceanic islands and seamounts are recognised as hotspots of abundance and diversity for marine top predators, such as sharks and large teleosts, and are hypothesised to function as migratory ‘stepping stones’ for mobile, pelagic species. However, while movements of pelagic predators between such features have been documented in some oceans, evidence from the tropical Atlantic remains limited.

**RESULTS:** Here, we report on long-distance migrations of three species of pelagic predator between oceanic islands and seamounts in the tropical south Atlantic recorded via independent passive acoustic telemetry arrays. These include the first recorded trans-Atlantic migration of a Galapagos shark (*Carcharhinus galapagensis*) from Ascension Island (UK) to the São Pedro and São Paul Archipelago (Brazil) (minimum distance 1,930 km); the longest documented oceanic migration of an Almaco jack (*Seriola rivoliana*) from St Helena (UK) to Ascension Island (minimum 1,300 km); and multiple movements of Galapagos and silky sharks (*C. falciformis*) between two Mid-Atlantic Ridge seamounts and Ascension Island (minimum 265–325 km). These detections are notable given the limited duration, sample sizes, and temporal overlap of acoustic tracking studies in the region, suggesting substantial connectivity across large spatial scales.

**CONCLUSIONS:** Our findings provide empirical support for the role of oceanic islands and seamounts as connectivity hubs for pelagic predators in the tropical Atlantic, underscoring their importance as key nodes in marine protected area networks. More broadly, this study demonstrates the value of collaborative regional tracking networks in resolving large-scale movement patterns and informing marine management at ecologically relevant scales.

## BACKGROUND

Continuing global declines in populations of many marine apex predators have created a pressing need to better understand patterns of distribution and movement in these species [1,2]. Many pelagic predators, including sharks, tuna, and billfishes are capable of traveling vast distances, routinely traversing entire ocean basins [1]. These long-distance migratory movements are of considerable ecological significance, linking distant populations, facilitating gene flow, and mediating energy transfer across otherwise disparate marine environments. Knowledge of migratory connectivity is also vital from a conservation perspective, informing fisheries management and supporting the design of well-integrated marine protected area (MPA) networks capable of accommodating the wide-ranging and often transboundary movements of these highly mobile species [1,2].

Seamounts and oceanic islands are widely recognised to be hotspots of abundance and diversity for a broad range of pelagic predators [3–5], functioning as both productive foraging ‘oases’ and aggregation areas for a variety of non-feeding related ecological processes (e.g. reproduction and refuging) [4]. According to the ‘stepping stone’ hypothesis, spatially connected chains of islands and seamounts may also play a key role in long-distance dispersal and migration, including acting as navigational aids or ‘refuelling stations’ for animals traversing expanses of comparatively unproductive and featureless open ocean [6]. Empirical support for such a role has predominantly been derived from satellite and acoustic tracking studies in the Pacific Ocean. For example, movements of multiple shark species have been documented among islands and seamounts in the Eastern Tropical Pacific [2,7,8], such as along the well-studied “Cocos-Galápagos Swimway” migratory corridor, while humpback whale migrations have been shown to track shallow topographic features in the South Pacific [6]. In comparison, evidence of connectivity between tropical Atlantic islands and seamounts is more limited. Weber et al. [4] recently reported movements of acoustically-tagged Galapagos sharks and silky sharks between two adjacent Mid-Atlantic Ridge seamounts (∼80 km), while Wright et al. [9] used a combination of conventional and satellite tags to record movements of yellowfin tuna (*Thunnus albacares*) between St Helena (UK) and two outlying seamounts (140–340 km). However, evidence of broader regional connectivity is lacking.

In this study, we synthesise data from several independent regional acoustic telemetry arrays across the equatorial and tropical south Atlantic to address this knowledge gap. Acoustic telemetry provides a powerful tool for studying pelagic predator movements among discrete topographic features, offering high positional accuracy, frequent transmissions, and long tag lifetimes (up to 10 years) [10]. These attributes enable detection of dispersal events that may be missed by shorter duration tracking methods with larger location error, such as satellite telemetry. However, its effectiveness depends on the simultaneous deployment of compatible receiver arrays and focused tagging of animals on multiple islands and seamounts over large spatial scales - an approach that has only recently become feasible in the tropical south Atlantic. With the expansion of acoustic telemetry infrastructure across the region, there is now a timely opportunity to assess connectivity of fully submerged aquatic species at broader spatial and temporal scales.

## METHODS

Acoustic telemetry data were compiled from four pelagic predator tracking studies that were conducted in the tropical Atlantic between 2017 and 2026 (Table 1). Study sites included Ascension Island (UK), the Grattan and Young Seamounts (UK), St Helena (UK), and the Saint Peter and Saint Paul Archipelago (SPSPA, Brazil). Ascension Island and St Helena are remote volcanic islands that administratively form part of a single UK Overseas Territory but are located approximately 1000 km apart in the South Atlantic Ocean (Fig 1.). The Grattan and Young Seamounts are a pair of shallow, mid-Atlantic Ridge features (<100 m depth) situated within Ascension Island’s exclusive economic zone, approximately 80 km apart and 260-320 km from the island [4]. SPSPA is a small group of rocky islets and seamounts located in the central equatorial Atlantic, ∼950 km from the Brazilian mainland and ∼90 km north of the equator. All sites are recognised hotspots for pelagic predators [4,11,12] and fall within MPAs designated since 2016 (Fig. 1).

**Table 1.**
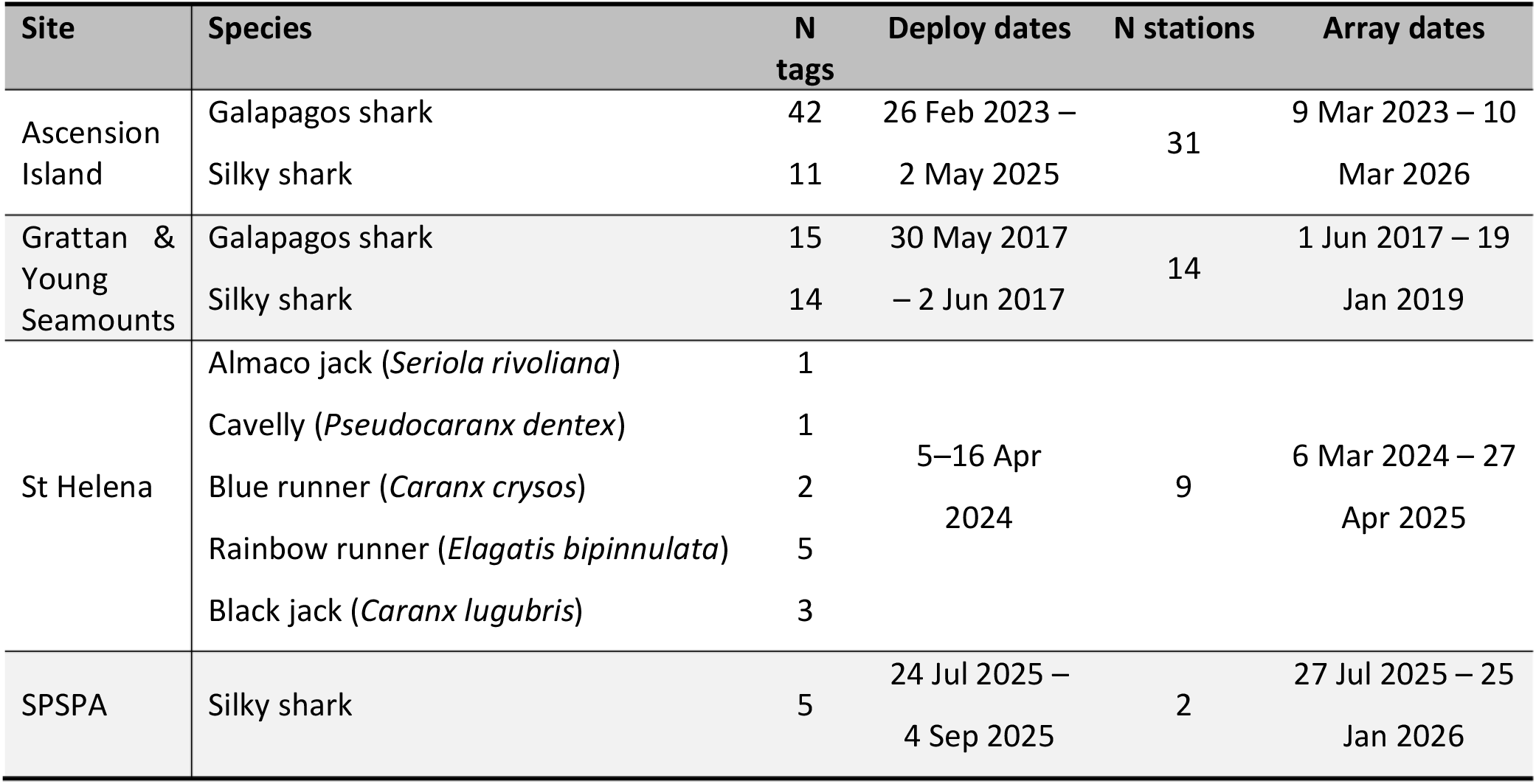
Summary of tropical Atlantic pelagic predator tagging studies included in the analysis. Total number of tags and receivers deployed at each sites are presented, along with timeframes for tag and acoustic tracking array deployments.

**Figure 1.**
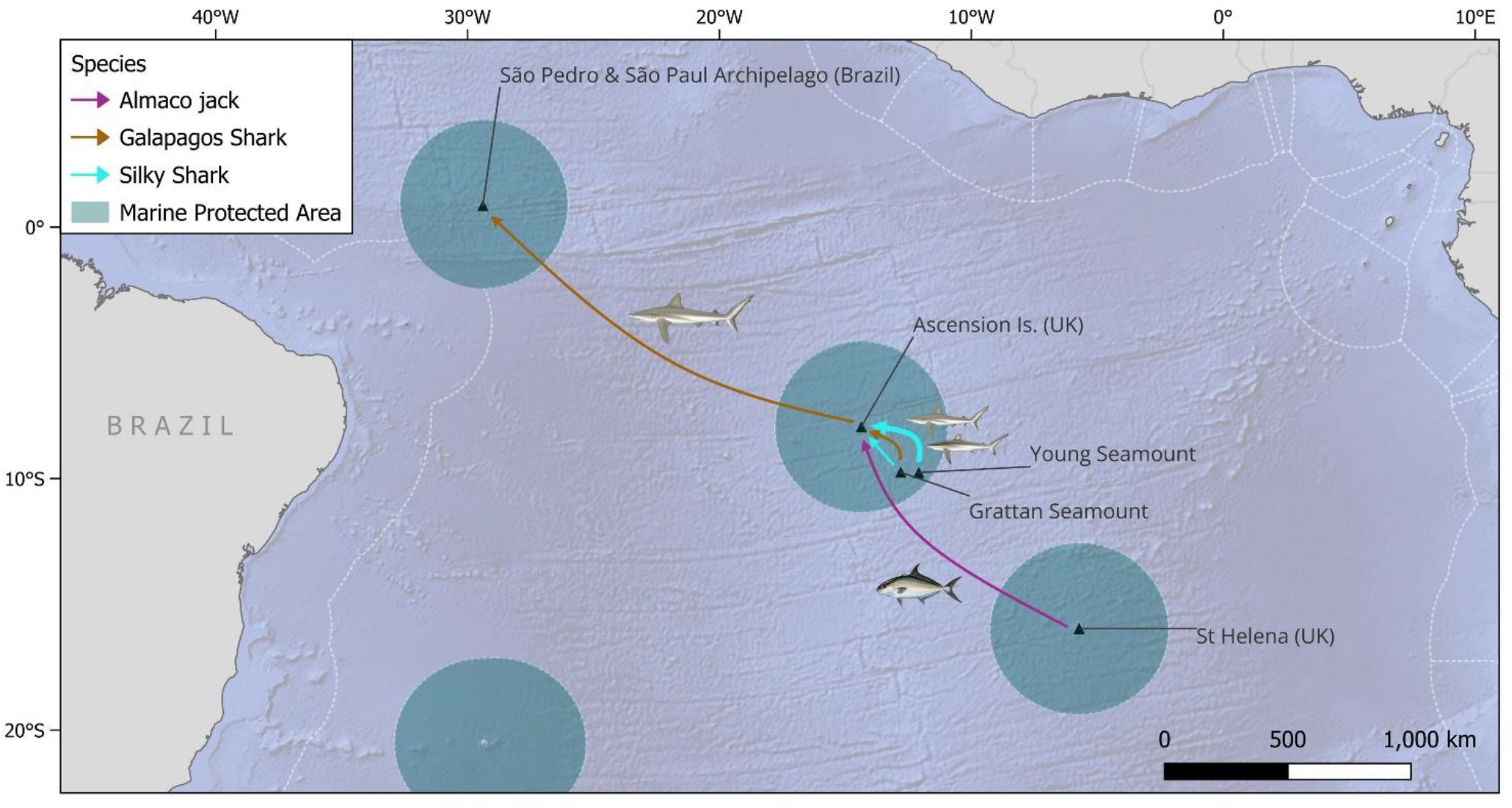
Long distance migrations of acoustically-tagged sharks and pelagic fish between islands and seamounts in the tropical south and equatorial Atlantic, recorded between 2023 and 2025. Basemap from www.naturalearthdata.com and MPA boundaries from www.protectedplanet.net. Illustrations: Sharks © Marc Dando; Fish © Diane Roam Peebles.

Acoustic tagging and installation of acoustic telemetry arrays at each site took place over different timescales with varying degrees of temporal overlap and with a range of different research objectives and focal species (Table 1; Fig. 2). At Ascension Island, two discrete tracking studies targeted Galapagos and silky sharks: one conducted at the Grattan and Young seamounts (2017–2018) and a second in coastal waters around Ascension Island itself (2023–2026), resulting in a total of 82 tagged individuals (Table 1). Recent work at SPSPA has similarly focussed on silky sharks, with five individuals tagged since 2025 (Table 1). In contrast, tagging at St Helena was more taxonomically broad with 12 individuals tagged across five species of predatory teleost (Table 1).

**Figure 2.**
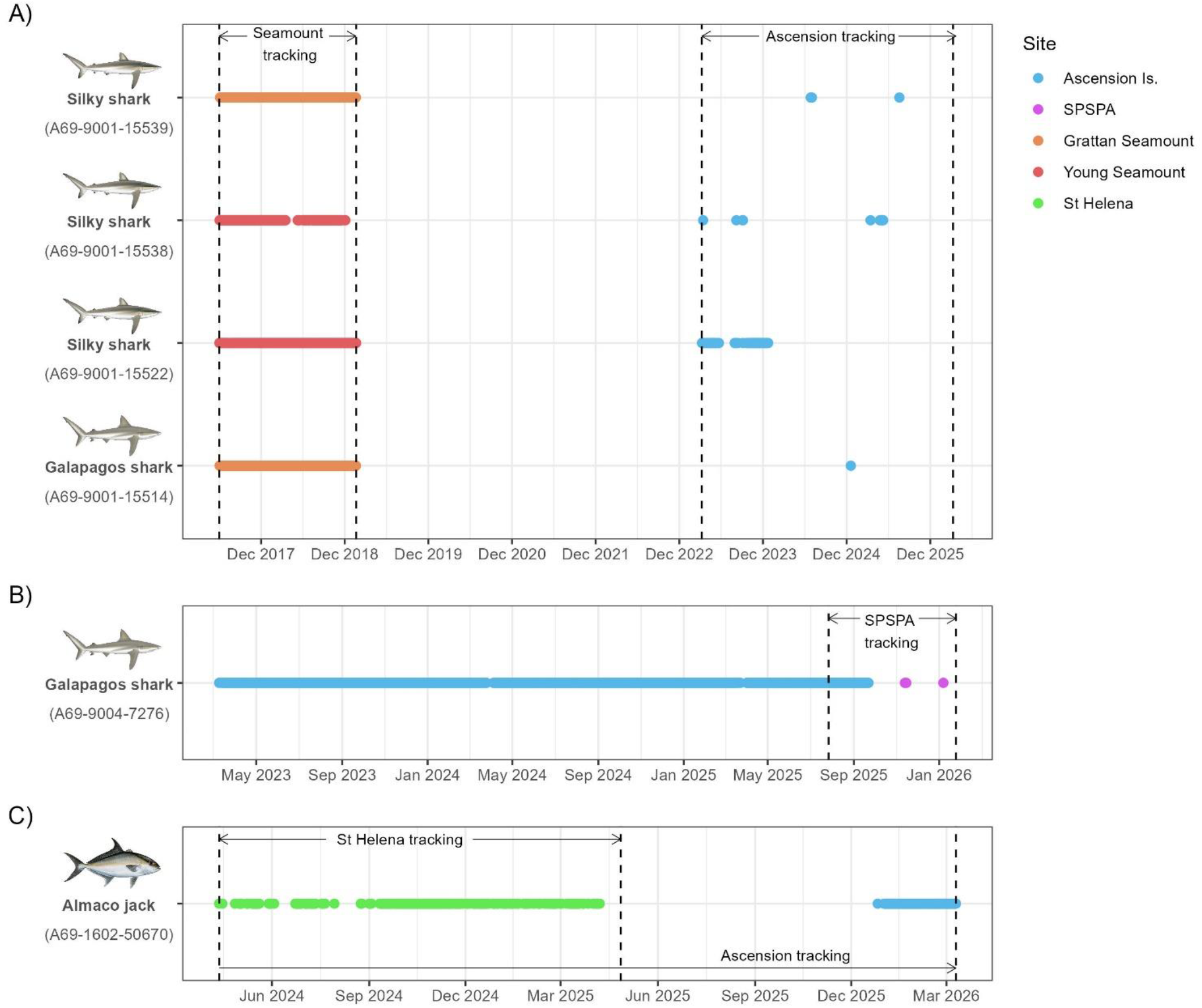
Abacus plot showing detections of acoustically-tagged pelagic predators that made long-distance migrations between islands and seamounts in the tropical Atlantic. Illustrations: Sharks © Marc Dando; Fish © Diane Roam Peebles.

At all sites, study animals were internally tagged with Innovasea 69 kHz acoustic transmitter tags (models V13 and V16), with estimated battery lives ranging from 6–10 years. All tagging was conducted under permits issued by relevant national authorities (Ascension: ERP-2023-001, ERP-2024-003, ERP-2025-003; SPSPA: SISBio No. 83121; St Helena: R12/2023) and using methods approved by institutional ethics committees (University of Exeter permit no. 845780; ZSL permit no. IOZ131; Universidade Federal Rural de Pernambuco permit no. CEUA-11564201123). Movements were then tracked using arrays of Innovasea 69 kHz receivers (models VR2-W, VR2-AR, and VR2-Tx) deployed in shallow waters around each feature (<150 m), with array sizes varying from two receivers at SPSPA to 31 at Ascension Island (Table 1). The use of standardised tag frequencies and compatible tag-receiver systems ensured that individuals were detectable across all sites within the regional network.

To explore connectivity between features, acoustic detection data from each site were screened for detections of individuals tagged elsewhere within the regional network. Detection validity was confirmed by Innovasea using a range of diagnostic filters to exclude false (‘ghost’) detections arising from acoustic signal collision, supported by evidence of repeated detections across multiple days and/or receivers in the array. For each confirmed inter-site movement, we estimated the minimum geodesic distance travelled and the maximum transit time, defined as the interval between the final detection at the departure site and the first detection at the arrival site. Residency at each site was quantified using both total residence times (time between first and last detections) and a residency index (RI), calculated as the proportion of days detected during the residency period. To estimate individual size at the time of inter-site movements (*L*_*t*_), we applied species-specific von Bertalanffy growth models with asymptotic size (*L*_∞_) and growth rate coefficients (*k*) obtained from published studies [13–15]:

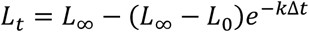

where *L*_0_ is the length at the time of tagging and Δ*t* is the time elapsed in years. Empirically derived allometric scaling relationships from the study sites were used to convert between alternative length measurements (pre-caudal, fork, and total length) where required. Estimates were derived using von Bertalanffy growth curves for male silky sharks from other Atlantic populations [13,14] and for Galapagos sharks in the Pacific [15] (since no Atlantic- or male-specific data were available).

## RESULTS

Between 2023 and 2025, a total of six long-distance migrations of acoustically-tagged sharks and teleost fish were detected between islands and seamounts in the central tropical Atlantic (Table 2 & Fig. 1). This included movements of three silky sharks and one Galapagos shark from the Grattan and Young Seamounts to Ascension Island (minimum distance = 265–325 km), a migration of a Galapagos shark from Ascension Island to SPSPA (minimum distance = 1,929 km), and a migration of an Almaco jack (*Seriola rivoliana*) from St Helena to Ascension (minimum distance = 1,300 km) (Figs. 1 & 2).

**Table 2.** Summary of individuals that made inter-site movements. AI: Ascension Island; SH: St Helena; GS: Grattan Seamount; YS: Young Seamount; SPSPA: São Pedro and São Paul Archipelago. *Fork length (FL) at the time of tag deployment.

| Species | Tag ID | FL (cm)* | Sex | Tag Date | Tagged at | Moved to |
| --- | --- | --- | --- | --- | --- | --- |
| Galapagos shark | A69-9004-7276 | 150 | M | 2023-02-28 | AI | ⇒ SPSPA |
| Almaco Jack | A69-1602-50670 | 61 | - | 2024-04-12 | SH | ⇒ AI |
| Galapagos shark | A69-9001-15514 | 135 | M | 2017-06-02 | GS | ⇒ AI |
| Silky shark | A69-9001-15538 | 143 | M | 2017-05-30 | YS | ⇒ AI |
| Silky shark | A69-9001-15522 | 121 | M | 2017-06-01 | YS | ⇒ AI |
| Silky shark | A69-9001-15539 | 135 | M | 2017-06-02 | GS | ⇒ AI |

All silky sharks and the single Galapagos shark that migrated from the Grattan and Young Seamounts to Ascension Island were males that were originally tagged as juveniles (121–143 cm FL; 161–180 cm TL) between May and June 2017 (Table 2). However, given the long interval between tag deployment and first detection at Ascension Island (5.2–7.4 years; Fig 2.), these individuals were estimated to have attained sizes >220 cm TL (>172 cm FL) by this stage. Since all individuals were still present at the Grattan and Young seamounts when the tracking study there ended in January 2019, the timing of migration to Ascension Island could not be determined precisely. However, two silky sharks were already present at Ascension when tracking there commenced in March 2023 (Fig. 2). Based on the elapsed time between tagging and final detections recorded at Grattan and Young (1.5–1.6 years), silky sharks were estimated to have been >=179–203 cm TL (139–161 cm FL) at the onset of migration, while the single Galapagos shark was estimated to have been >=192 cm TL (162 cm FL).

The Galapagos shark detected moving from Ascension Island to SPSPA was also a male that was originally tagged as a large juvenile or sub-adult (150 cm FL; Table 2). In this case, overlap between acoustic monitoring periods at both sites allowed the timing of movement to be more precisely constrained. The final detection of this individual at Ascension Island occurred on 22^nd^ September 2025, 937 days (2.6 years) after tagging, and it was first detected at SPSPA 52 days later, on 13^th^ November 2025. Estimated size at the onset of migration was 221 cm TL (188 cm FL). Based on estimated mean cruising speeds for a 188 cm FL carcharhinid shark (0.73–0.84 m/s [16]), the minimum direct transit time between Ascension and SPSPA is approximately 27–31 days, suggesting that this individual may have made more exploratory movements before arriving at SPSPA.

All Galapagos and silky sharks that undertook long distance migrations exhibited a similar pattern of residency around topographic features, characterised by an extended period of highly residential behaviour at the tagging site followed by more transient and infrequent detections at the features they migrated to (Fig. 2). Individuals tagged at the Grattan and Young Seamounts were almost continuously resident around these features over a 595-day period (RI = 0.84–1.0) but were detected only intermittently at Ascension Island after moving, with residence times ranging from 1–124 days and RI ranging from 0.01–0.43 (Fig 2). Two of the silky sharks were detected at Ascension on only 4 and 14 days respectively over a ∼3-year tracking period, with evidence of repeated visits after absences of ∼12–18 months, while the single Galapagos shark that migrated from the Grattan Seamount was detected on only a single day (39 detections on five different receivers). Similarly, the single Galapagos shark that migrated from Ascension Island to SPSPA remained strongly associated with Ascension for >2.6 years post-tagging (RI = 0.95) but was detected on only three days at SPSPA over a 55-day period (RI = 0.05) (Fig. 2)

In contrast, the Almaco jack that migrated from St Helena to Ascension, exhibited extended periods of residency at both locations. Following tagging, this individual remained highly resident around St Helena for approximately 1 year (RI = 0.55) until acoustic monitoring there ceased in April 2025. It was first detected on Ascension Island in December 2025 where it remained for the subsequent 74 days until tracking ceased in March 2026, generating more than 2,500 detections across 18 receivers (RI = 0.95). The prolonged residency and repeated movements of this fish at Ascension Island make it unlikely that the tag had been secondarily transported following predation, as reported tag retention times after ingestion are typically in the order of days to weeks e.g. [17]. Because receiver deployments at St Helena and Ascension Island did not fully overlap temporally, the timing of movement could not be determined precisely, although it occurred between 360 and 620 days after tagging. By this stage, the individual is expected to have grown substantially beyond its tagging size of 61 cm FL; however, in the absence of species-specific growth parameters, it was not possible to estimate length at the time of movement.

## DISCUSSION

Our results provide empirical evidence that oceanic islands and seamounts function as connectivity hubs for pelagic predators in the tropical south Atlantic, consistent with patterns documented in other ocean basins [2,7,8]. Across all detected movements, sharks exhibited extended periods of high residency at their tagging locations prior to undertaking long-distance dispersal events, followed by comparatively transient residency at the sites to which they migrated. This pattern suggests that oceanic islands and seamounts may function both as long-term aggregation habitats and as intermittently used “stepping stones” that facilitate broader-scale movement across the open ocean. Notably, all recorded shark movements involved males and, based on estimated body size, occurred around the onset of sexual maturity, which is attained at a mean size of 200–220 cm TL in silky sharks [13,14] and 200–240 cm TL in Galapagos sharks [18]. While this apparent male bias may partly reflect sampling biases in the tagged populations (which were >72% male at all sites), sex-biased dispersal associated with maturation has been documented in a range of shark species [19] and may represent an important driver of connectivity between habitats. The comparatively transient detections observed following arrival at new sites may therefore partly reflect ontogenetic shifts in habitat use, whereby individuals become less strongly associated with shallow island or seamount habitats after reaching adulthood.

The most frequent movements detected were those of Galapagos and silky sharks from the Grattan and Young Seamounts to Ascension Island (a minimum distance of 260–320 km). In total, three of 14 silky sharks (20%) and one of 15 Galapagos sharks (6%) tagged as juveniles on the seamounts were subsequently detected at Ascension Island, despite a 6-year gap between tagging and the initiation of acoustic monitoring at Ascension. This is comparable to the proportion of individuals reported to make shorter distance (∼80 km) movements between Grattan and Young Seamounts (silky: 36%; Galapagos 13%; [4]), indicating a continuum of habitat connectivity among topographic features within the Ascension Island Marine Protected Area.

The more frequent long-distance movements observed in silky sharks are consistent with the ecology of this species in other regions. Although silky shark distributions are often strongly associated with topographic features, particularly during juvenile stages [4,20,21], they are widely recognised as highly mobile pelagic predators capable of extensive oceanic migrations [21,22]. By comparison, Galapagos sharks are generally considered a more coastal pelagic species and often exhibit strong site fidelity to oceanic islands, seamounts, and shelf-edge habitats [4,8,23]. Nevertheless, more wide-ranging movements and occasional long-distance dispersal between topographic features have previously been documented in the Eastern Tropical Pacific [8,24,25], including a reported >3000 km migration between the Revillagigedo Archipelago, Clipperton Atoll, and the Galápagos Islands [8]. To our knowledge, the >1,900 km movement of a Galapagos shark from Ascension Island to the São Pedro and São Paul Archipelago (SPSPA) reported here represents the first confirmed large-scale migration of this species in the Atlantic Ocean and demonstrates that similar trans-oceanic dispersal events also occur in this basin.

The long-distance (>1,300 km) migration of an Almaco jack from St Helena to Ascension Island additionally demonstrates that connectivity among topographic features in the tropical Atlantic is not restricted to sharks, but may also occur in large predatory teleosts. Although Almaco jack are reported to be highly resident around seamounts and oceanic islands [26], as also observed here, population genetic studies have reported limited genetic structuring within ocean basins, including the Atlantic, consistent with a degree of connectivity among populations [27]. However, long-distance oceanic migrations have not been previously documented in this species.

Taken together, our results support the hypothesis that oceanic islands and seamounts may function as key connectivity hubs, genetically and demographically linking pelagic predator populations across broad spatial scales. If corroborated by additional data, these preliminary findings have important implications for the conservation of pelagic predators in the tropical Atlantic, as demographic trends at one site may influence connected populations elsewhere, contributing to regional population dynamics and resilience. For example, the Galapagos shark population at SPSPA was once considered functionally extinct following decades of commercial fishing pressure [28] yet appears to have rebounded rapidly since legal protection in 2012 [29]. While this recovery may reflect intrinsic population growth, our results suggest that immigration from more stable regional populations, such as those at Ascension Island, may also have contributed to population replenishment. This underscores the importance of MPA networks that reflect ecological connectivity among geographically separated habitats.

Notably, all movements documented in this study occurred between large-scale pelagic marine protected areas (LSMPA) established within the past decade. While LSMPAs have attracted criticism for failing to adequately encompass the migratory ranges of highly mobile pelagic species (reviewed in [30]), our results suggest that protecting strategically important connectivity hubs, such as islands and seamounts, may nonetheless yield substantial conservation benefits for some feature-associated species by safeguarding key aggregation and movement nodes. Indeed, evidence of similar connectivity in the Eastern Tropical Pacific recently informed the development of the world’s first transboundary marine reserve [2], highlighting the value of comparable datasets from other oceans to inform regional-scale marine management. Although relatively few inter-site movements were recorded in this analysis, they are notable given the sparse receiver coverage across the tropical Atlantic, incomplete temporal overlap between studies, and small sample sizes, with only five features monitored over varying timescales and some species represented by only a single individual. Based on these preliminary findings, we anticipate that the expansion of a more coordinated, spatially comprehensive, and long-term acoustic tracking network across the tropical Atlantic, modelled on similar initiatives elsewhere (e.g. [31]), would likely reveal substantially higher levels of regional connectivity. Such data would be highly valuable in resolving large-scale movement dynamics of pelagic predators in the Atlantic and informing the design of well-connected MPA networks capable of addressing ongoing declines in many of these species.

## ACKNOWLEDGEMENTS

Work on Ascension Island was funded by the UK Darwin Initiative (grant no. DPLUS063 & DPLUS161), the EU BEST Initiative (grant. no. 1599), a Save Our Seas Foundation Keystone Grant (SOSF700), and the UK Government’s Conflict Stability and Security Fund. Work on St Helena was funded by St Helena Government and the UK Government’s Blue Belt Programme. The Ocean Tracking Network provided acoustic telemetry equipment used in the SPSPA study. The authors are grateful for the support of all members and partner institutions of the BRISCA Network and current and former Ascension Island Government and St Helena Government employees and volunteers who assisted in fieldwork and data management, including: Andy Richardson, Judith Brown and Cuen Muller (Ascension Island); Adam Riggs, Cerys Joshua, Peter Benjamin, Rico Benjamin, and Clayton Benjamin (St Helena). We would also like to thank the operators and crews of the multiple vessels that supported fieldwork logistics, including the M/V Extractor, RRS James Clarke Ross, AIG Marine Operations, Craig Yon (Dive St Helena) and Anthony Thomas (Subtropic Adventures), the Brazilian Navy, ICMBio Grandes Oceânicas, and Lunus.

